## Supplementary figures for "Systems approaches identify the consequences of monosomy in somatic human cells"

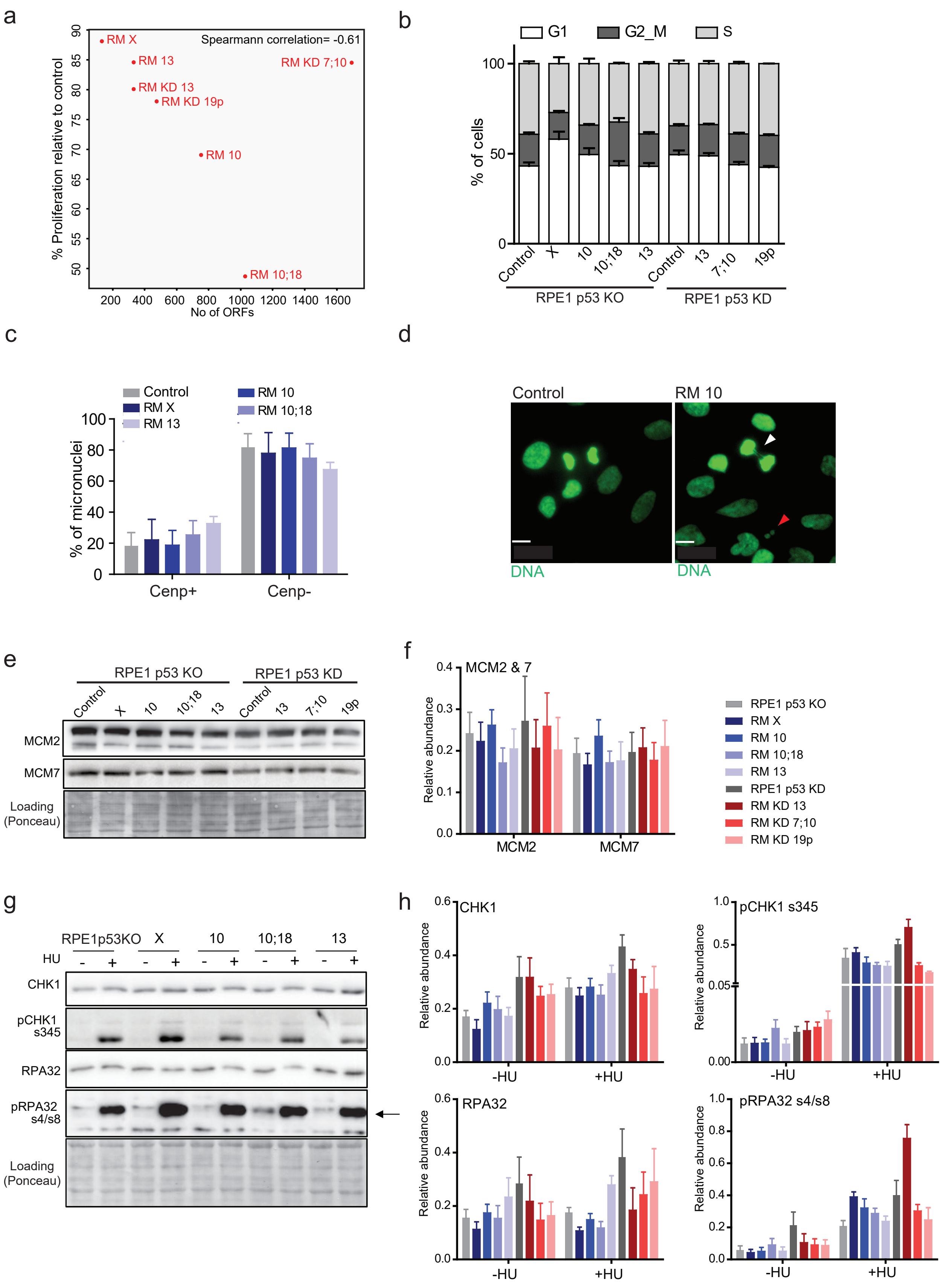


Chunduri et al

Supplementary figure 1 **Characterization of monosomic cell lines**

**a**. Correlation between the proliferation and number of open reading frames (ORFs) on the monosomes. The number of ORFs was obtained from NCBI database; the estimated number of non-compensated ORFs was considered for chromosome X. **b**. Cell cycle profiles of control and monosomic cell lines. **c**. Fraction of micronuclei positive/negative for CENP signal. **d**. Representative image of anaphase bridges in diploid and monosomic cell lines. Scale bar – 10 µm. **e**. Representative immunoblot of the subunits of the key replicative helicase MCM2-7. **g**. Quantification of the protein abundance from f. **h**. Representative immunoblot of key DNA damage response factors and **i**. Quantification of their abundance. Arrow points to the specific band for pRPA32 s4/s8. All plots depict mean with SEM, at least three independent biological experiments were performed. HU-hydroxyurea.

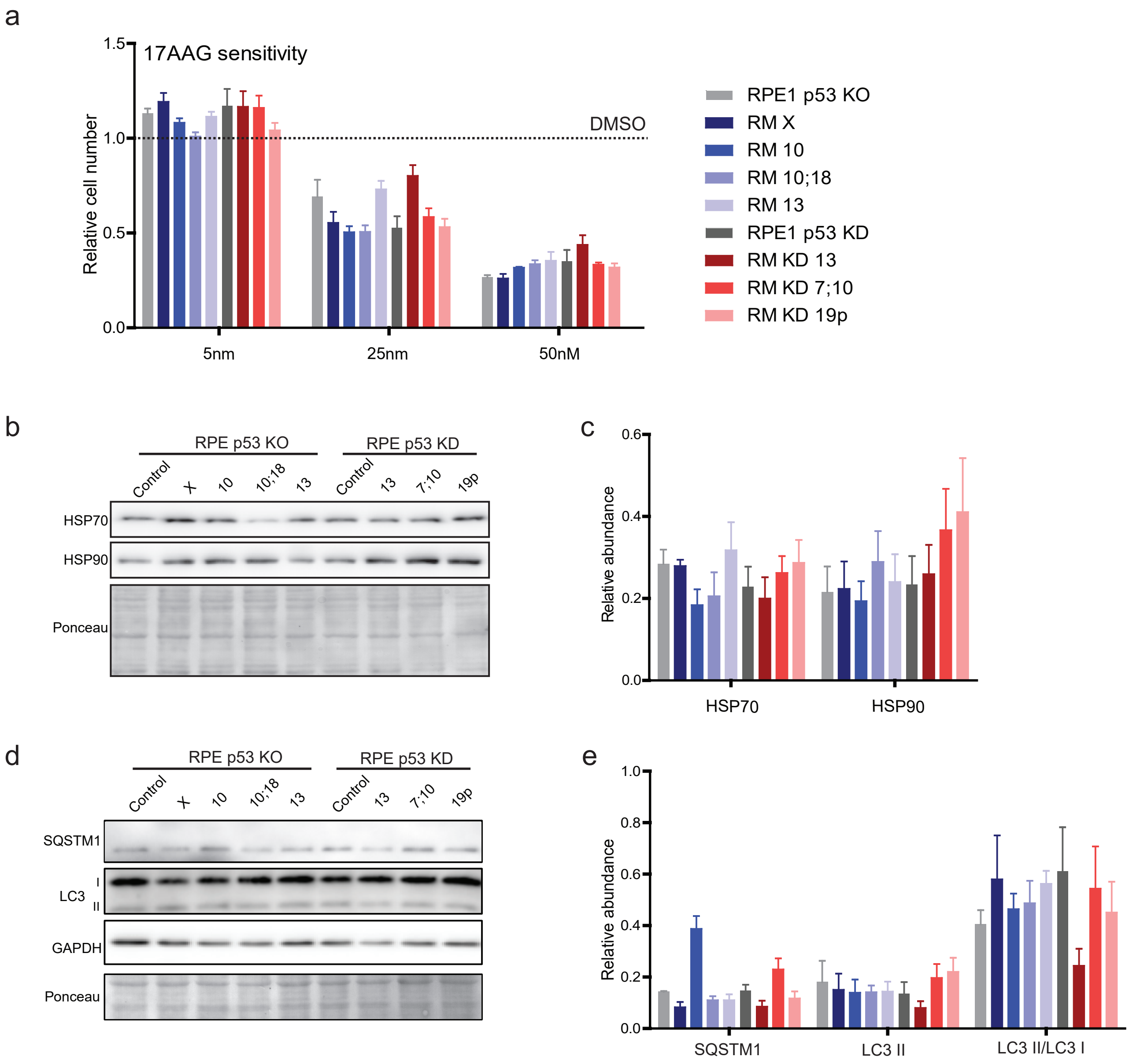


Chunduri et al

Supplementary figure 2 **No apparent proteotoxic stress in monosomies.**

**a.** Sensitivity to 17AAG measured by Cell Titer Glo assay. All the values were normalized to DMSO control.
**b.** Immunoblotting of heat shock proteins and **c.** quantifications of at least three independent experiments.
**d.** Immunoblotting of autophagy related proteins and **e.** quantification of at least three independent experiments.
Mean and SEM is shown in all plots. Ponceau staining was used as a loading control in (**b**) and (**d**).

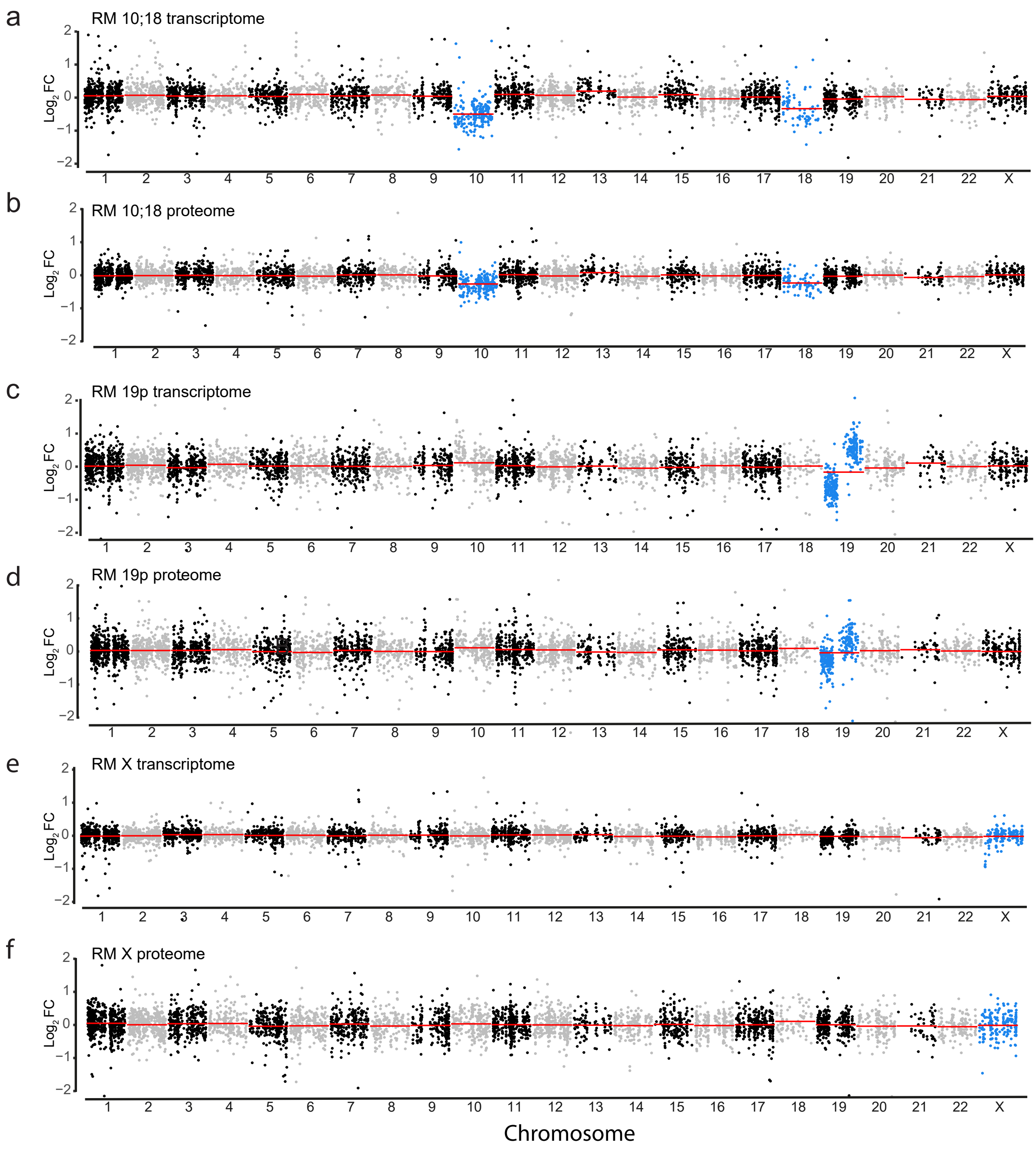


Chunduri et al

Supplementary figure 3 **Transcriptome and proteome of monosomic cell lines.**

**a-f.** The relative abundance of mRNAs and proteins of RM10;18, RM 19p and RM X normalized to diploid control and plotted according to the chromosome location of the corresponding genes. The monosomic chromosomes are marked in blue. Red line depicts the median for each chromosome.

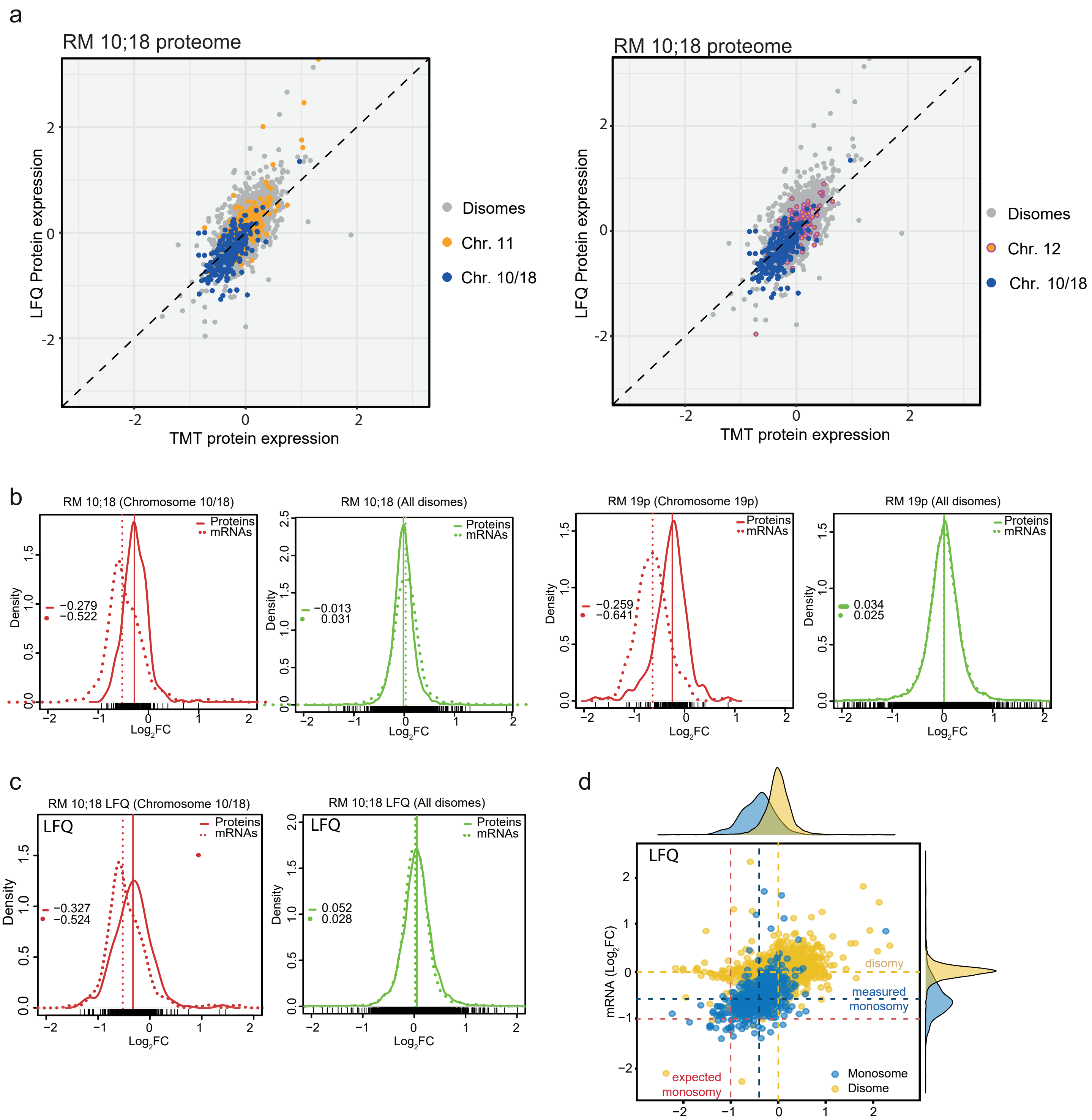


Chunduri et al

Supplementary figure 4. **Comparative values were obtained by LFQ and TMT proteomics analysis.**

**a.** Scatter plots of the relative protein abundance values obtained by LFQ and TMT proteomics show a strong similarity between the results obtained by these two approaches. RM10;18 proteome is shown, proteins encoded on chromosomes 10 and 18 (monosomic) and 11 and 12 (disomic) are highlighted. All other disomic chromosomes are shown in light gray. **b.** Overlay of mRNA and protein density histograms for RM 10;18 and RM 19p. Red line depicts the monosomes, green line the disomes. **c.** Overlay of mRNA and protein density histograms for RM 10;18 using the LFQ data. **d.** Scatter plot showing the log2 fold change (FC) of mRNA and proteins encoded on monosomes (blue) and disomes (yellow), as in Fig. 3d, but calculated based on the LFQ data. The marginal density histograms show the distribution of respective mRNAs and proteins. The expected median fold change of monosomic genes is marked by red dashed lines. The measured median fold changes of monosomic and disomic genes is marked by blue and yellow dashed lines, respectively.

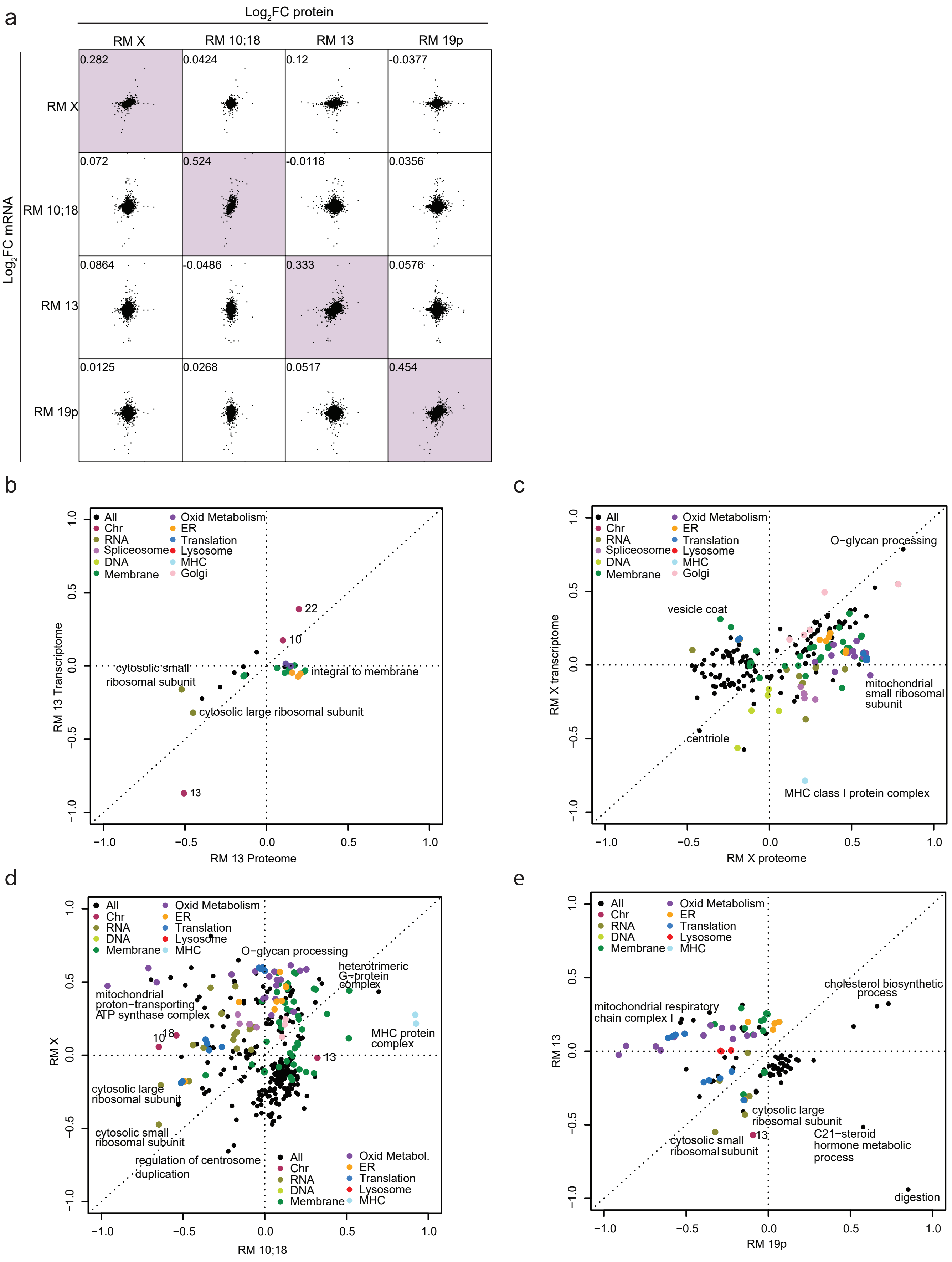


Chunduri et al

Supplementary figure 5 **Pathway enrichment analysis of monosomies**

**a.** Scatter plots depicting the Spearmann rank correlation coefficient between the relative abudances of mRNA and proteins for all monosomies. The numbers in each plot represent the Spearmann correlation coefficient. **b,c**. 2D pathway enrichment analysis of transcriptome and proteome of RM 10;18 and RM X. **d,e**. 2D pathway enrichment analysis comparing the proteome of individual monosomic cell lines.

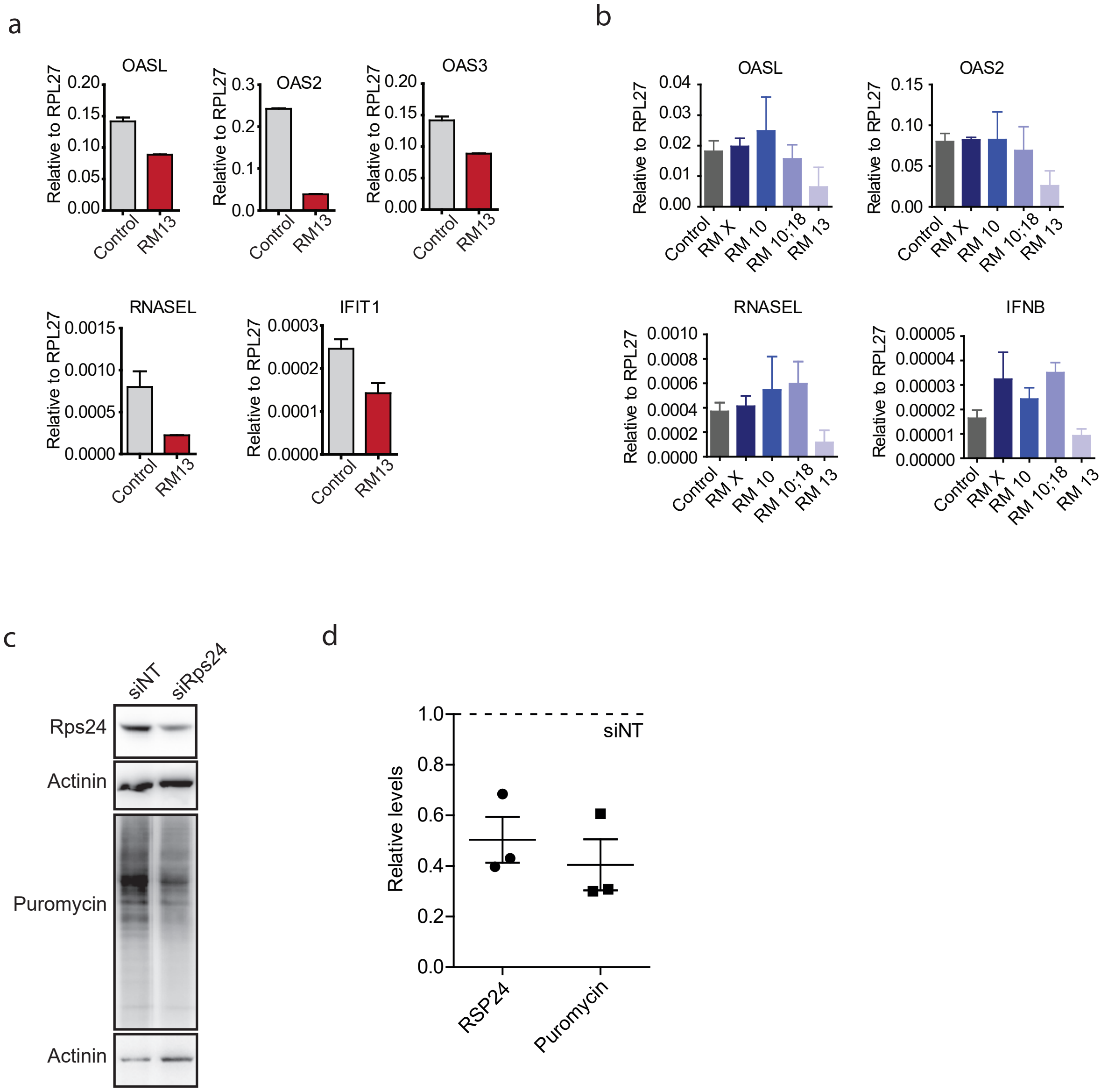


Chunduri et al

Supplementary figure 6 **Distinct and shared expression changes in response to monosomy**
**a, b.** Monosomic and control cells were treated with 100U of Interferon B and the expression of the interferon reponse genes was quantified by qPCR. **c** Representative blot of RPS24 levels upon siRNA for RPS24 and for non-targeting control (NT). Levels of RPS24 and puromycin incorporation are shown. $\alpha$– actinin was used as a positive control. **d**. Quantification of three independent experiments as in c. The expression of RPS24 and Puromycin in RPS24 transfected cells was normalized to NT control. Mean with SEM is shown.

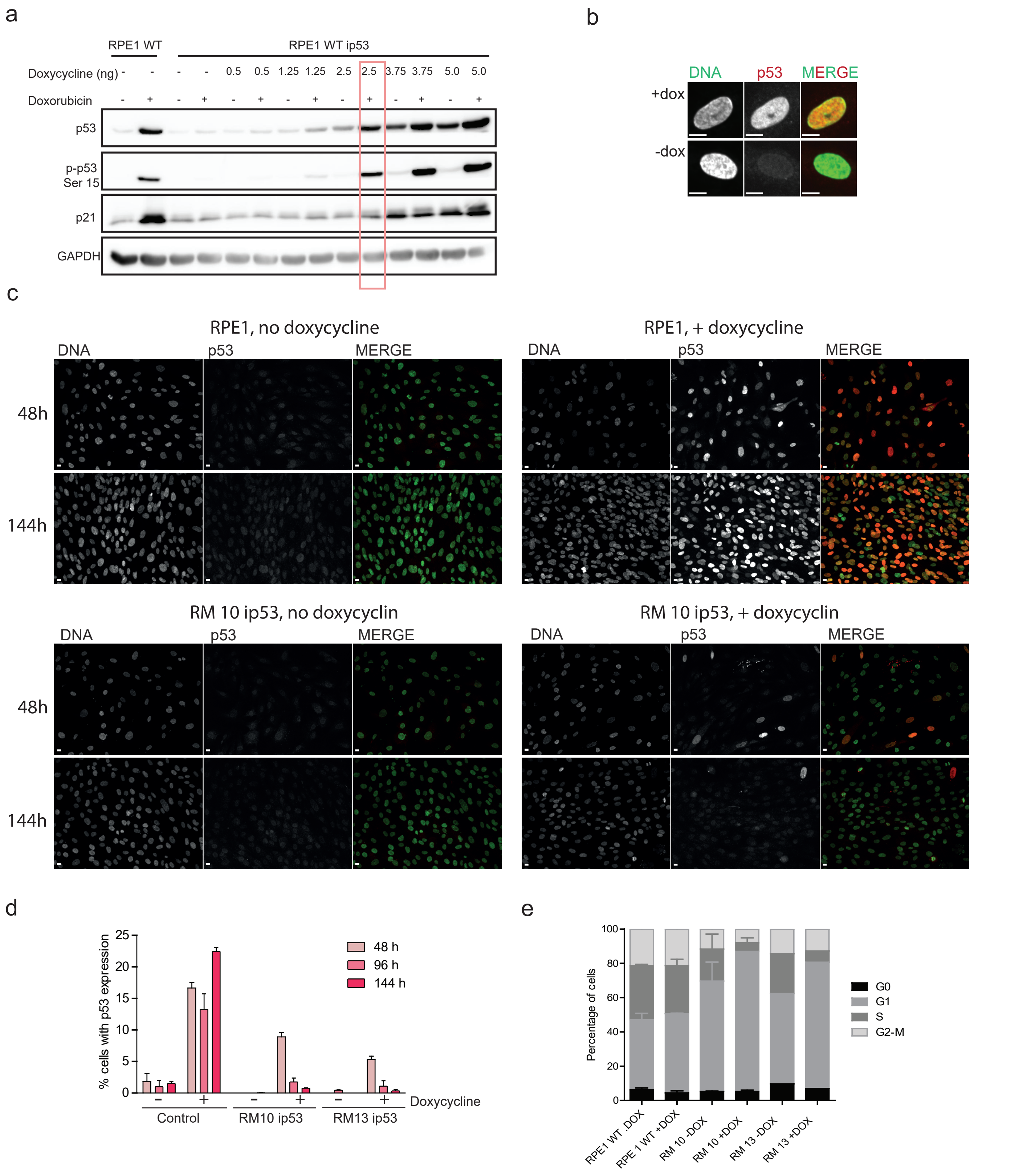


Chunduri et al

Supplementary figure 7 **Restoration of p53 in monosomic cell lines**

**a.** Titration of the doxycycline to reach normal p53 expression in RPE1 WT ip53 cell line. RPE1 WT is used as a control. Cells were treated with either doxycycline (in nanograms) or doxorubicin, or both. GAPDH serves as a loading control. Marked levels (red box) were used for the experiments. **b, c.** Immunofluorescence staining of p53 in cells with and without doxycycline treatment. DNA was stained with Sytox green, p53 is visualized in red. Scale bar - 10 µm. **d.** Quantification shows the percentage of p53 expressing cells with and without treatment with doxycycline. Bars display mean ± SEM of at least two independent experiments. **e.** Cell cycle profile of monosomies with and without p53. Percentage of cells in different cell cycle phases were plotted.

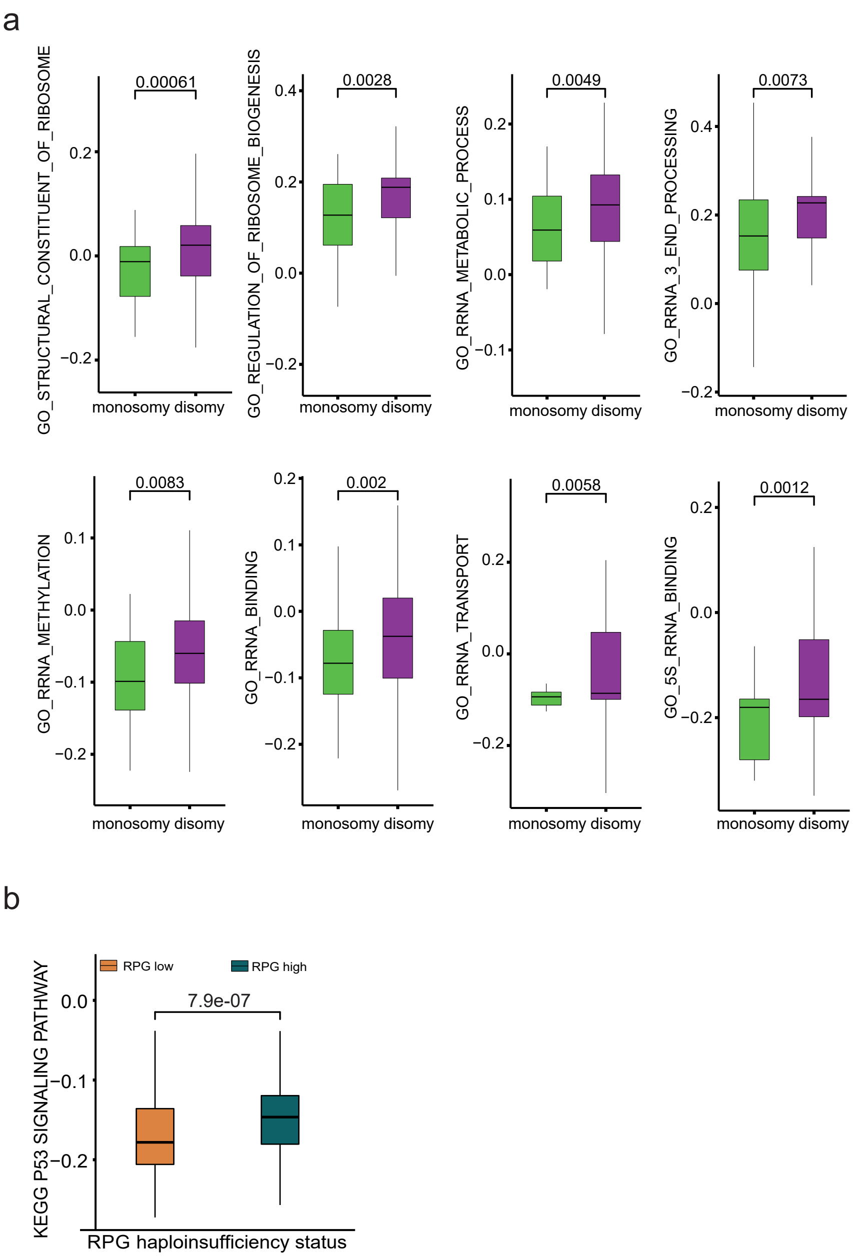


Chunduri et al

Supplementary figure 8 **CCLE transcriptome data comparing monosomy cell lines to disomy cell lines**

**a.** Transcriptomic analysis shows the ssGSEA enrichment score of GO terms related to ribosomes and rRNA in monosomic cell lines compared to disomic cell lines. **b.** KEGG p53 pathway score of monosomy cell lines from CCLE. Monosomy cell lines were divided into two groups based on the median RPG abundance of the entire cohort. Orange box denotes group whose RPG expression is lower than the median of the cohort and blue box represents the group with RPG expression higher than the median of the cohort. Wilcoxon rank sum text for used to evaluate the statistical significance in all figures, p-values are shown in the respective plots.
