## Supplementary table 1 for "Systems approaches identify the consequences of monosomy in somatic human cells"

Supplementary table 1 Engineered monosomic cell lines

| **Name** | **Parent** | **Altered chromosome** | **Fraction of monosomic cells** | **Remarks** |
| --- | --- | --- | --- | --- |
| RM10 | RPE1-hTERT p53 -/- | 10 | Chr.10  100% (N=25) | This work |
| RM 10;18 | RPE1-hTERT p53 -/- | 10 and 18 | Chr.10  100% (N=20) | This work |
| RM X | RPE1-hTERT p53 -/- | X | Chr. X  100% (N=15) | This work |
| RM 13 | RPE1-hTERT p53 -/- | 13 | Chr. 13  91% (N=11) | This work |
| RM13 kd | RPE1-hTERT shRNA-p53; H2B-Dendra2 | 13 | N/A | Soto et al, 2017 |
| RM 7;10 kd | RPE1-hTERT shRNA-p53; H2B-Dendra2 | 7 and 10 | N/A | Soto et al, 2017 |
| RM 19p kd | RPE1-hTERT shRNA-p53; H2B-Dendra2 | 19p | N/A | Soto et al, 2017 |
| RPE1 ip53 | RPE1-hTERT p53 -/- |  | 2N | This work |
| RM 13 ip53 | RM 13 | 13 | Chr.13  85% (N=13) | This work |
| RM 10 ip53 | RM 10;18 | 10 and 18 | Chr.10  75% (N=24) | This work |

Fraction of monosomic cells:

Percentage of metaphase spreads having one copy of respective monosomic chromosome and 2 copies of diploid chromosome.

N= number of metaphase spreads
