## Supplementary table 6 for "Systems approaches identify the consequences of monosomy in somatic human cells"

| **Antibodies** | | |
| --- | --- | --- |
| **Name of the protein** | **Company** | **Identification number** |
| P53 (DO-1) | Santa Cruz | Sc-126 |
| Anti puromycin 12D10 | Merck Millipore | MABE343 |
| p21 Waf1/Kip1 | Cell signaling | 2947 |
| p-eIF2 alpha (Ser51) | Cell signaling | 9721S |
| eIF2 alpha | Cell signaling | 9722S |
| LC 3a/b | Cell signaling | 4108 |
| p70 S6 Kinase | Cell signaling | 2708 |
| p-p70 S6 Kinase | Cell signaling | 9205 |
| Ribosomal Protein L21 (D7) | Santa Cruz | Sc-393663 |
| alpha-actinin | Santa Cruz | sc-17829 |
| HSP90 | Cell signalling | 4874 |
| HSP70/HSP72 | Enzo | ADI-SPA-902 |
| Chk1 | Abcam | Ab32531-100 |
| p-Chk1 | Cell signalling | 2348 |
| CDK2 | Cell signalling | 2546 |
| CDK6 | Cell signalling | 3136 |
| Cyclin D1 | Cell signalling | 2978 |
| Cyclin B1 | Cell signalling | 4135 |
| P18 | Cell singalling | 2896 |
| MCM2 | Abcam | Ab4461 |
| MCM7 | Santa Cruz | Sc9966 |
| pRPA32(s33) | Bethyl | A300-246A |
| pRPA32(s4/s8) | Bethyl | IHC-00422 |
| RPA32 | Abcam | Ab2175 |
| P62 lck ligand | BD Transduction | 610832 |
| Cenp B | Santacruz | Sc376392 |
| γH2AX | Abcam | Ab2893 |
| **siRNA** | | |
| siRPL21 | Dharmacon | M-012910-01-0005 (Smartpool)  Sequences:  GUACCUGGGUUCAACUAAA  GAGAAUUAAUGUGCGUAUU  GAGGAGAGGCACCCGAUAU  CCACAUAUAUGCGAAUCUA |
| siGENOME Non-Targeting Control siRNA Pool #1 | Dharmacon | D-001210-01-05(Smartpool)  Sequences:  UAGCGACUAAACACAUCAA  UAAGGCUAUGAAGAGAUAC  AUGUAUUGGCCUGUAUUAG  AUGAACGUGAAUUGCUCAA |
